# Ambient temperature crystal structure of *Escherichia coli* CyaY protein displays alternate conformation

**DOI:** 10.1101/2023.03.08.531761

**Authors:** Alaleh Shafiei, Nilufer Baldir, Jongbum Na, Jin Hae Kim, Hasan DeMirci

**Affiliations:** Department of Molecular Biology and Genetics, Koc University, 34450, Istanbul, Turkey; (A.S.); (N.B.); (H.D.); Department of New Biology, Daegu Gyeongbuk Institute of Science and Technology (DGIST), 42988, Daegu, Republic of Korea; (J.N.); (J.H.K.); Koc University Isbank Center for Infectious Diseases (KUISCID), Koc University, 34010, Istanbul, Turkey; Stanford PULSE Institute, SLAC National Laboratory, Menlo Park, CA 94025, USA, (H.D.)

**Keywords:** CyaY, Frataxin, Friedreich ataxia, *Escherichia coli*, Turkish Light Source *“Turkish DeLight”*, X-ray crystallography, Ambient-temperature crystallography, Structural biology

## Abstract

Frataxin is a 23 KDa mitochondrial iron-binding protein that is involved in biogenesis of iron sulfur clusters. A deficiency in frataxin leads to Friedreich’s ataxia, a progressive neurodegenerative disorder. The bacterial ortholog of eukaryotic mitochondrial frataxin, CyaY, is thought to play a role in iron sulfur cluster assembly as an iron supplier, making it an important target for study. Here, we present the first ambient temperature crystal structure of CyaY protein from *Escherichia coli*, obtained using the Turkish Light Source “Turkish DeLight”. This study reveals the dynamic structural characteristics of CyaY at near-physiological temperature and displays an alternate conformation, highlighting the importance of temperature considerations in protein structure characterization and providing new insights into the protein’s flexibility.

## 1. Introduction

Iron sulfur clusters (ISCs) have been identified as one of the most ancient redox-active co-factors in the cell [1]. These clusters have been found to play a critical role in a diverse range of biological processes, including electron transport and transfer, gene expression, gene regulation, photosynthesis, nitrogen fixation, enzyme-substrate binding, DNA repair and replication, and RNA modification [2,3,4,5,6]. The assembly of iron sulfur clusters is accomplished through universally conserved systems present in prokaryotic, archaeal and eukaryotic worlds [7], These systems have the potential to provide Fe-S clusters to a wide range of apo-proteins [8,9]. In prokaryotes, three major classes of systems responsible for the formation of iron-sulfur clusters have been identified: i) NIF (Nitrogen fixation) system [10], ii) SUF (sulfur mobilization) system and iii) ISC (iron sulfur cluster) system, each responsible for the formation of iron sulfur clusters in housekeeping proteins in stress and normal conditions, respectively [11,12,13]. These distinct systems share a common feature in the utilization of the cysteine desulfurase enzyme (NifS, SufSE, IscS) to obtain sulfur atoms from cysteine[14], which then conjugate with an iron atom to form an iron sulfur cluster on a scaffold protein (NifU, SufB, IscU) or cluster precursor intermediates [15,16,17].

The *Escherichia coli* Iron sulfur cluster system is the most well-studied as a model system because of its lower complexity and it encodes several protein units that have eukaryotic homologs. These include IscR (a transcriptional regulator) [18,19], IscS (a cysteine desulfurase) [20,21], IscU (a scaffold protein which clusters are built on) [22], ferredoxin (an electron supplier) [23], a chaperone/co-chaperone system which then helps to transfer the cluster from IscU to the target apo-proteins (HscA, HscB) [24], IscX (a potential iron supplier) [25]. Cluster formation needs the intricate interactions and interplays of these multi proteins and other accessory proteins whose roles are not well understood yet. One such partner is the frataxin protein. In eukaryotes frataxin deficiency results in loss of iron sulfur cluster protein activity and is associated with Friedreich’s ataxia, an autosomal recessive neurodegenerative disease in humans [26,27]. The frataxin has been proposed to function as an iron-donor, an iron-chaperone or iron sensor regulator of the iron sulfur cluster system [28,29,30]. CyaY is of interest to researchers as it is the bacterial ortholog of eukaryotic mitochondrial frataxin [31,32]. Moreover, it exhibits remarkable evolutionary conservation from bacteria to higher eukaryotes [31] and shares 25% sequence identity with human and yeast frataxin (FXN/Yfh1) [32]. Biochemical research has demonstrated that in the presence of IscS, CyaY can donate an iron and can mediate formation of iron sulfur clusters on IscU *in vitro* [32]. Furthermore, the binary complex of IscS-CyaY was identified by isothermal titration calorimetry (ITC) and small-angle X-ray scattering (SAXS) studies which revealed that IscS and CyaY bind each other [33]. Additionally, ternary complexes of IscS-IscU-CyaY were also characterized by pulldown assays [34] and cross linking studies [35]. Therefore, it is believed that there is a link between CyaY as an iron donor/transporter and the biogenesis of iron sulfur clusters.

In this study, we aimed to gain a deeper understanding of the macromolecular structural dynamics of the bacterial ortholog of frataxin, CyaY from E. coli, by determining its first ambient temperature structure. Prior structural studies on CyaY and biogenesis of iron sulfur clusters have been conducted in cryogenic temperature [36,37,38,39]. Despite the insights they provided, as a consequence of the fact that proteins freeze in the intermediate timescales while cryocooling [40], lowered temperatures have the potential to limit the identification of alternative protein conformations that might be functionally important [41]. In addition, they have failed to show structural and functional heterogeneity of the complexes involved in the cluster biogenesis so far. To address these limitations, we determined the crystal structure of CyaY protein from *E. coli* at ambient temperature by using our *Turkish DeLight* home X-ray source. The X-ray crystal structure presented through this experiment, therefore, may lay the foundation for further studies in understanding the underpinnings of the mechanism of iron sulfur cluster assembly machinery in near-physiological temperatures. These findings may also contribute to a deeper understanding of the biogenesis mechanisms of Fe-S clusters in mitochondria and the development of therapeutic strategies for neurodegenerative diseases associated with disruptions in these processes.

## 2. Materials and Methods

### 2.1. Protein Expression and Purification

The N-terminal hexahistidine-Sumo-tagged CyaY gene was transformed and overexpressed in *E. coli* Rosetta2™ BL21 (DE3) strain. The cells were grown at 37° C in 4L LB media supplemented by 50 μl/ml kanamydn and 35 μl/ml chloramphenicol for large scale protein expression until OD_600_ reached the 0.6-0.8 range. Then 1:1000 (v:v) 0.4 mM Isopropyl β-D-1-thiogalactopyranoside (IPTG) was added at 0.4 μM final concentration and overexpression continued at 18 °C for 18h. Cells were harvested at 3500 rpm by Beckman Allegra 15R desktop centrifuge and resuspended in a lysis buffer containing 20 mM Tris–HCl (pH 7.5), 500 mM NaCl, 30 mM imidazole, and 5 mM β-mercaptoethanol. Then cells were lysed by sonication by using Branson W250 sonifier (Brookfield, CT, USA) followed by ultracentrifugation at 35000 rpm at Beckman Optima™ L-80XP ultracentrifuge equipped with Ti-45 rotor to isolate the soluble supernatant extract. The affinity purification was performed using Ni-NTA affinity column (Qiagen, Venlo, Netherlands), and the purified CyaY protein was then placed in the dialysis buffer and treated by Ulp1 to remove the N-terminal hexahistidine-Sumo tag. Cleaved CyaY was separated from uncleaved protein by passing the protein mixture through the Ni-NTA affinity column, and the flowthrough, containing pure cleaved CyaY protein, was collected. The cleaved native CyaY protein was further concentrated to a final concentration of 3 mg/ml using (Amicon 10KDa MWCO) ultrafiltration columns and it was kept in −80 for future experimentation.

### 2.2. Crystallization

Sitting drop vapor diffusion microbatch under oil crystallization technique was used for crystallization of CyaY protein at 4 °C. Search for crystallization conditions was performed by adding purified CyaY to commercial sparse matrix and grid crystal screen conditions in the 72 well Terasaki crystallization plates with the 0.83 μl protein to cocktail ratio of 1:1 (v/v). In total approximately 3000 commercial sparse matrix and grid screen crystallization conditions were set up and examined [42]. CyaY crystals were obtained in Grid-NaCl crystallization conditions containing 0.1 M Citric acid and 4.0 M Sodium chloride (pH 4.0) (Hampton Research, USA) within two weeks.

### 2.3. Ambient temperature Data Collection and Processing

Ambient temperature X-ray crystallographic data was collected using Rigaku’s XtaLAB Synergy R Flow XRD system, as outlined in Gul *et.al*. 2022 [43]. The diffraction data was then merged and processed with CrysAlisPro 1.171.42.59a software (Rigaku OxfordDiffraction,2022), as described in Atalay *et al* 2022 [44].

### 2.4. Structure Determination and Refinement

The phase problem was solved using molecular replacement with PHASER-MR in the PHENIX *suite* [45], using the cryogenic CyaY crystal structure (PDB code: 1EW4) as the starting model [45]. After simulated-annealing refinement, individual coordinates and TLS parameters were refined. Potential positions of altered side chains and water molecules were checked and the models were built/rebuilt by using the program *COOT* [46]. Final structure refinement was performed using phenix.refinement in *PHENIX* [46]. Structural figures were generated using *PyMOL* (Schrödinger, LLC)(https://pymol.org/). Data processing and refinement statistics are listed in (Table 1).

**Table 1.**
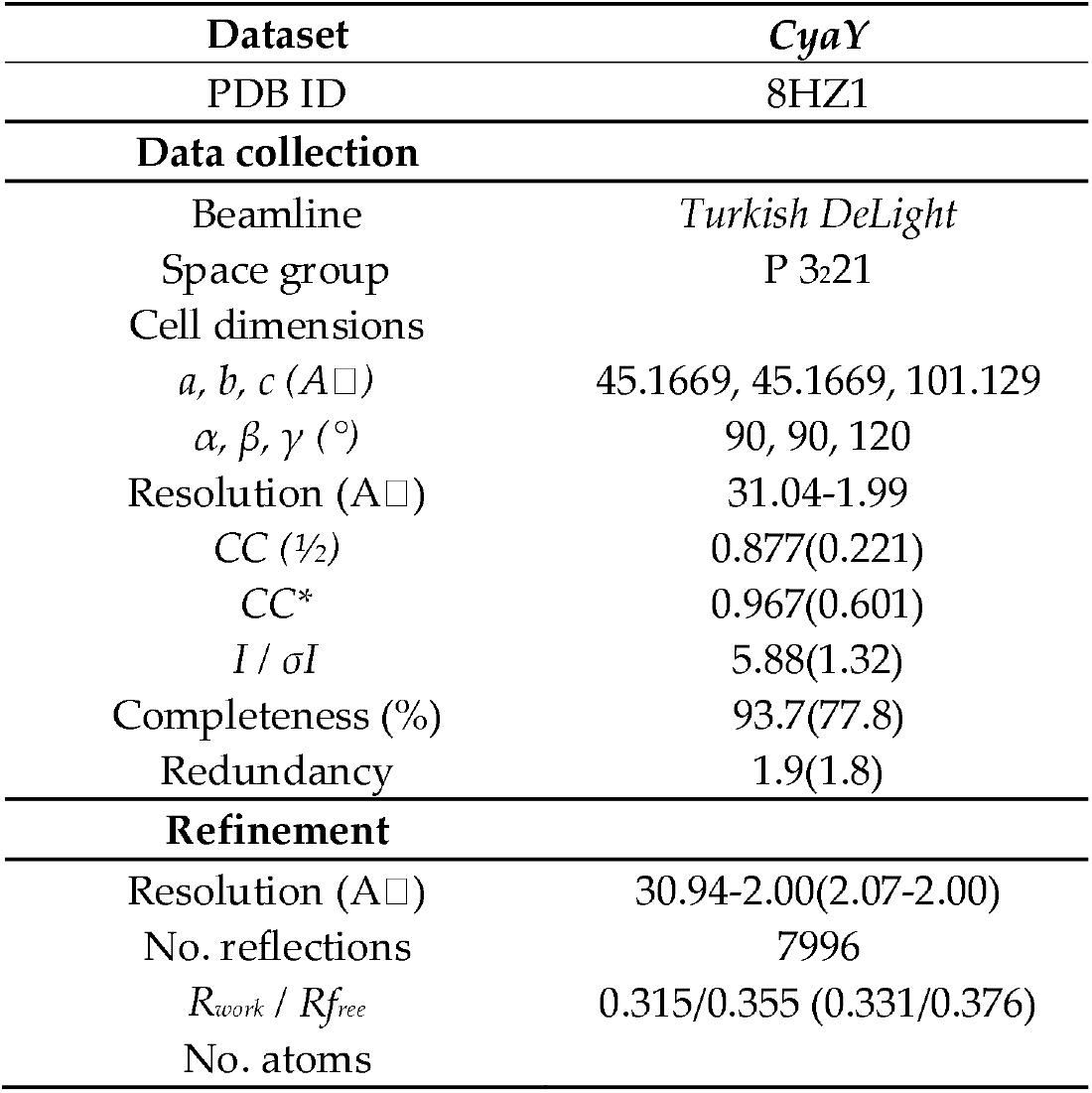
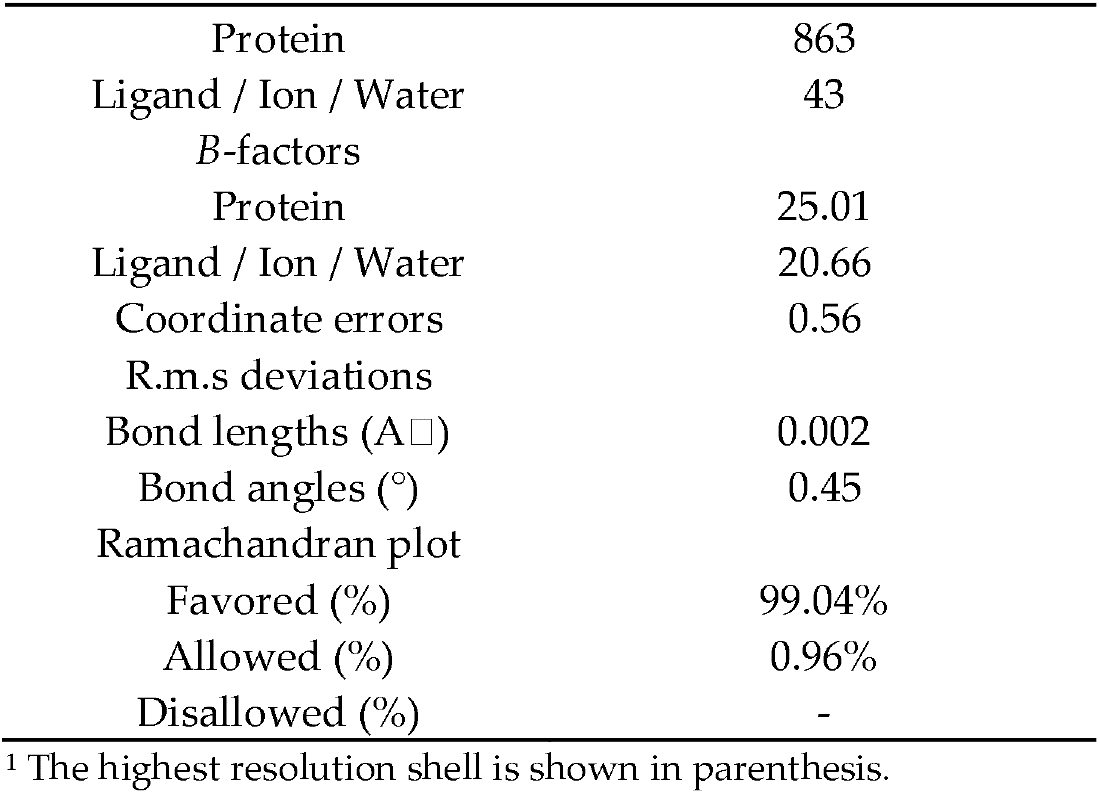
Data collection and refinement statistics.

## 3. Results

### 3.1. Ambient temperature CyaY structure

Ambient temperature crystal structure of CyaY protein was determined (PDB ID: 8HZ1) in its monomeric form at a resolution of 2 Å in space group P3_2_21 (Figure 1 and Table 1). The structure comprises 106 residues consisting of 3 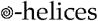 and 6 β-strands (Figure 1). Analysis of Ramachandran plot showed that 99.04% of residues are within the favored regions, while 0.96% of residues are in the allowed regions. The structure contains no Ramachandran outliers. The monomeric structure of CyaY is composed of a twisted antiparallel β-sheet, which is surrounded by 2 *α*-helices. The beta-sheet is formed by six adjacent beta-strands (β1-β6) and is enclosed by two long alpha-helices (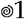 and 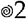) at the N-terminus and C-terminus, respectively. Additionally β3 and β4 are connected by a short helix called 3_10_ (Figure 1). Details Structural information and refined electron density maps (2Fo-Fc) for the residues are shown in (Figure 1).

**Figure 1.**
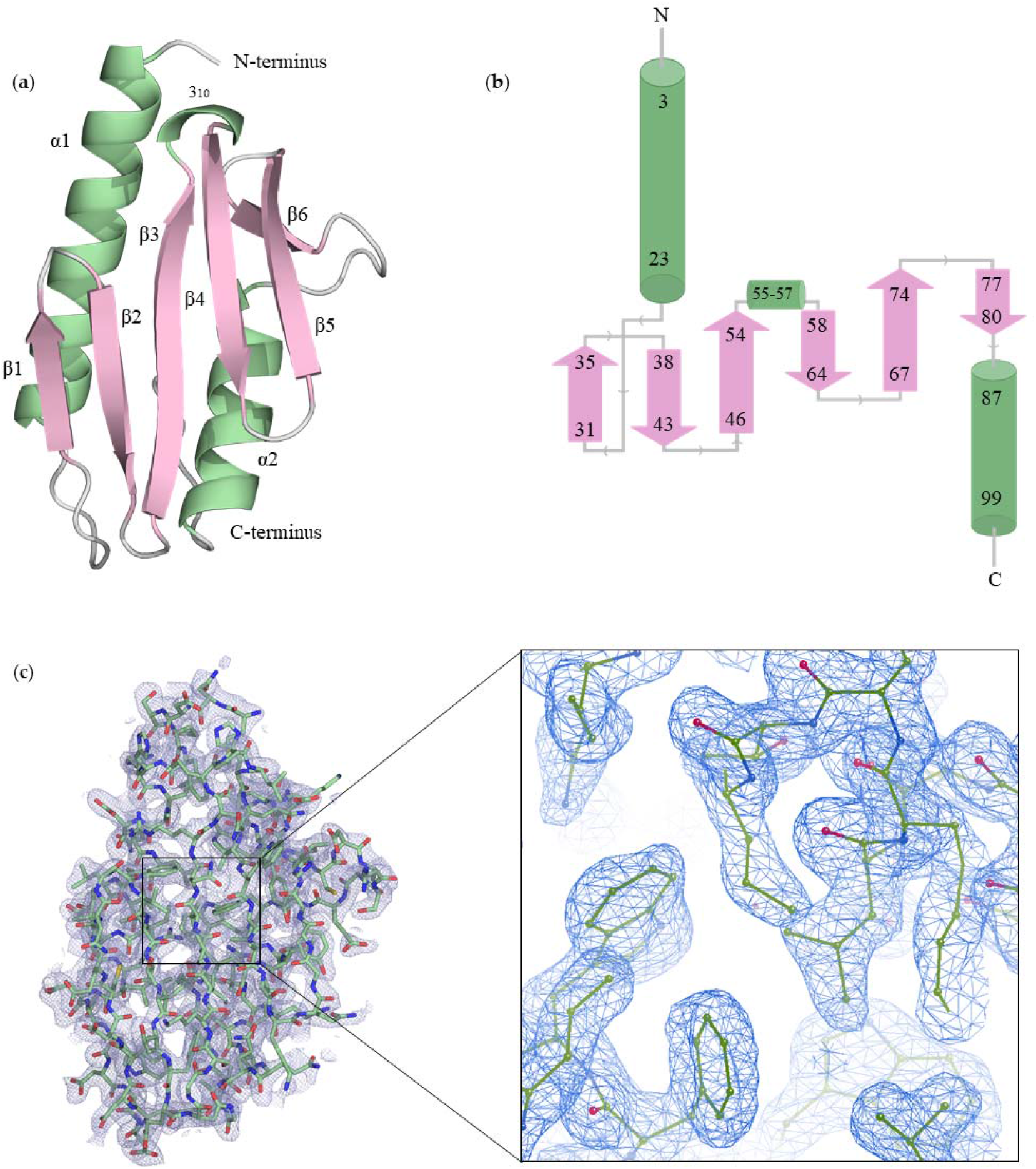
(a) 2 Å resolution monomeric crystal structure of *E. coli* CyaY protein. Alpha helices, Beta strands and loops are shown in deeptail, pink and pale green, respectively, (b) Topology map. Pink arrows indicate beta strands, green cylinders represent alpha helices, and lines between shapes indicate loops. (c) 2Fo-Fc map is colored in blue and contoured at 1.0 σ level. CyaY is shown in stick representation here.

### 3.2. Comparison with Cryogenic structure

A superimposition of the ambient temperature structure with the cryogenic structure (PDB ID:1EW4) revealed that the β strands in the cryogenic structure (β2, β3, β4, β5, β6) are shorter in length when compared to our ambient temperature structure (Figure 2). This suggests the presence of small conformational changes in the ambient temperature structure. Further details regarding the β strands can be found in (Figures 3, 4, & 5). The analysis also revealed minor conformational changes in residues Ile79 and Glu5, while significant changes were observed in residues Asn2, Arg8, Asp23, Asp76, Leu90, and Gln66 (Figure 6).

**Figure 2.**
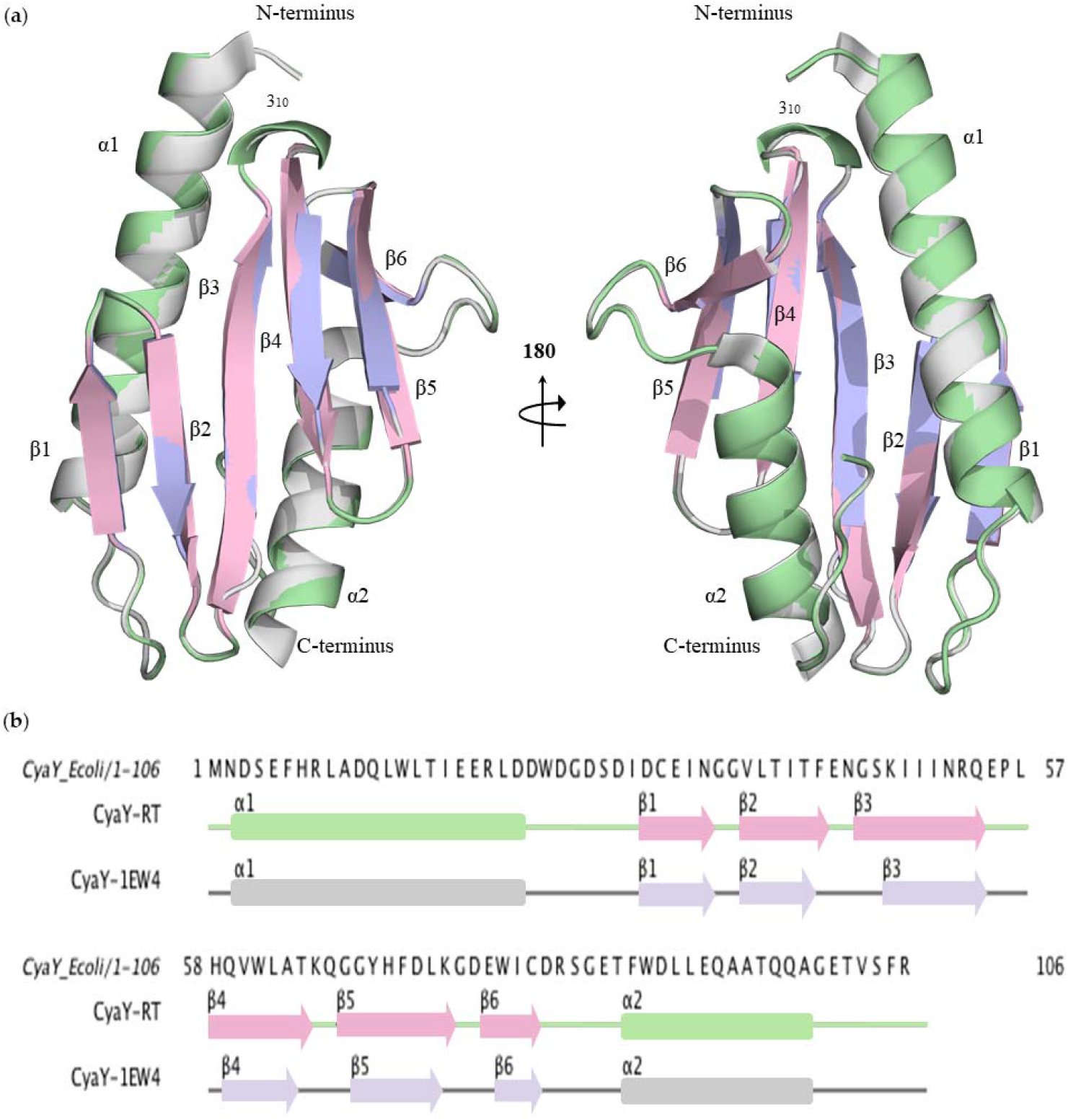
Comparison of secondary structures of CyaYs. **(a)** Superimposition of CyaY structure at ambient temperature with the cryogenic structure (PDB:1EW4) with RMSD value of 0.194. Two side views are presented in the panel by rotating the structure 180 degrees on the y-axis. **(b)** Structure-based sequence alignment of CyaY at ambient temperature with the cryogenic structure. Generated by *JALVIEW*. Alpha helices are depicted in deeptail and light blue, Beta sheets are shown in pink and gray, Loops are illustrated in pale green and gray for RT(room temperature) and cryogenic temperatures, respectively.

**Figure 3.**
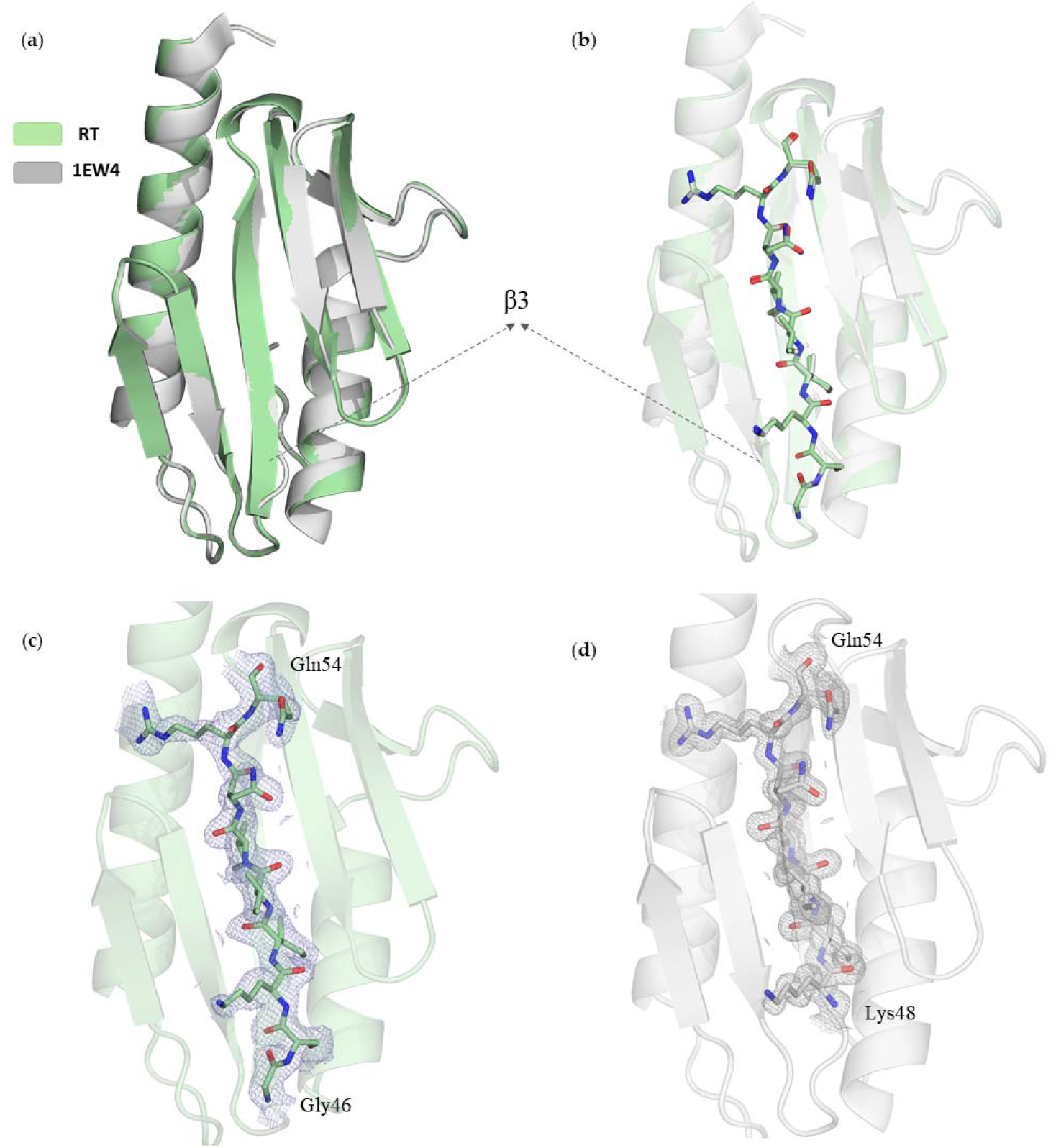
Comparison of β3 strand, **(a-b)** Superimposition of RT and cryogenic temperature CyaY protein. **(c-d)** stick representation of β3 strand in RT and cryogenic structure, respectively.

**Figure 4.**
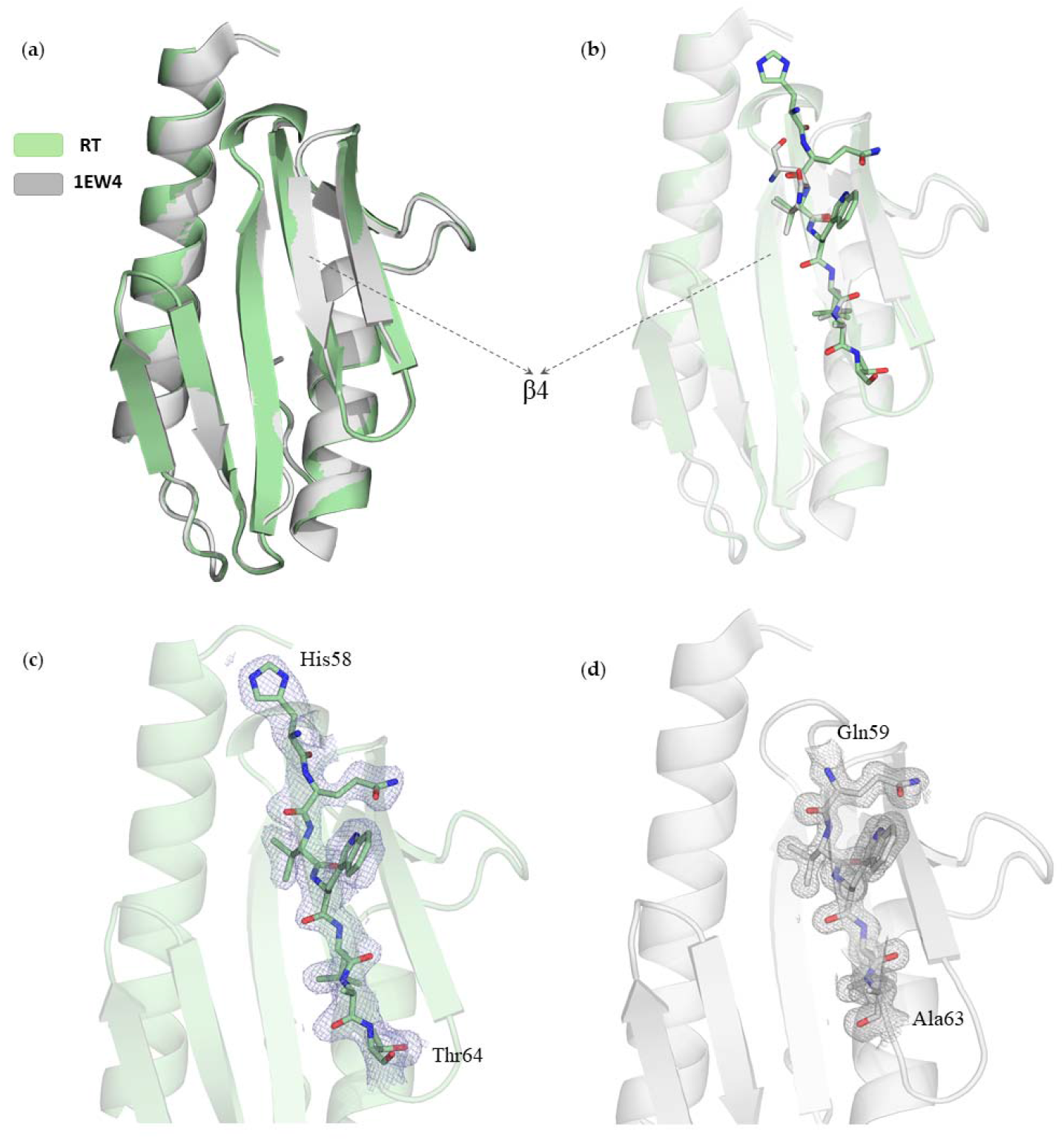
Comparison of β4 strand, **(a-b)** Superimposition of RT and cryogenic temperature CyaY protein. **(c-d)** Stick representation of β4 strand in RT and cryogenic structure, respectively.

**Figure 5.**
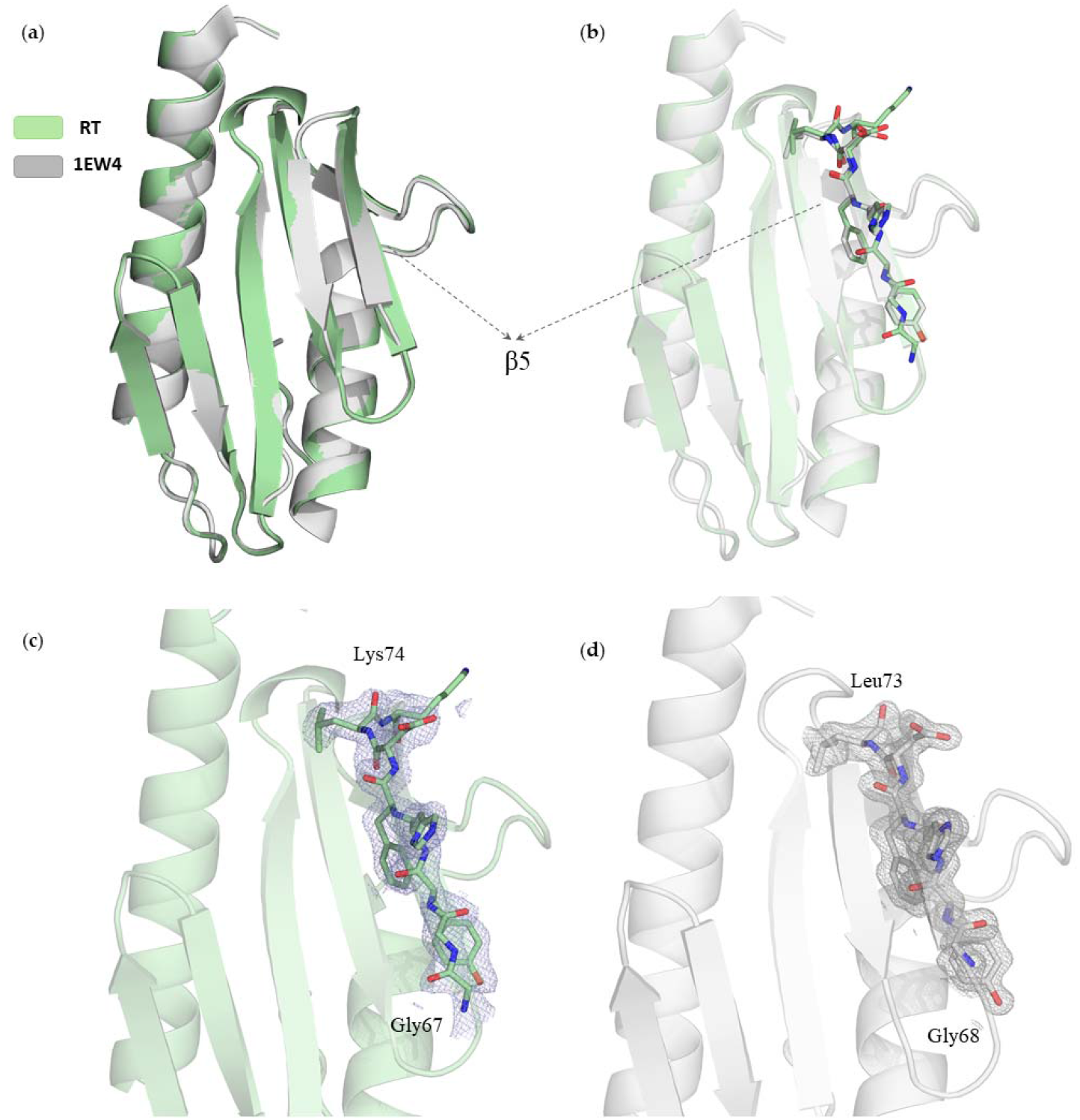
Comparison of β5 strand, **(a-b)** Superimposition of RT and cryogenic temperature CyaY protein. **(c-d)** stick representation of β5 strand in RT and cryogenic structure, respectively.

**Figure 6.**
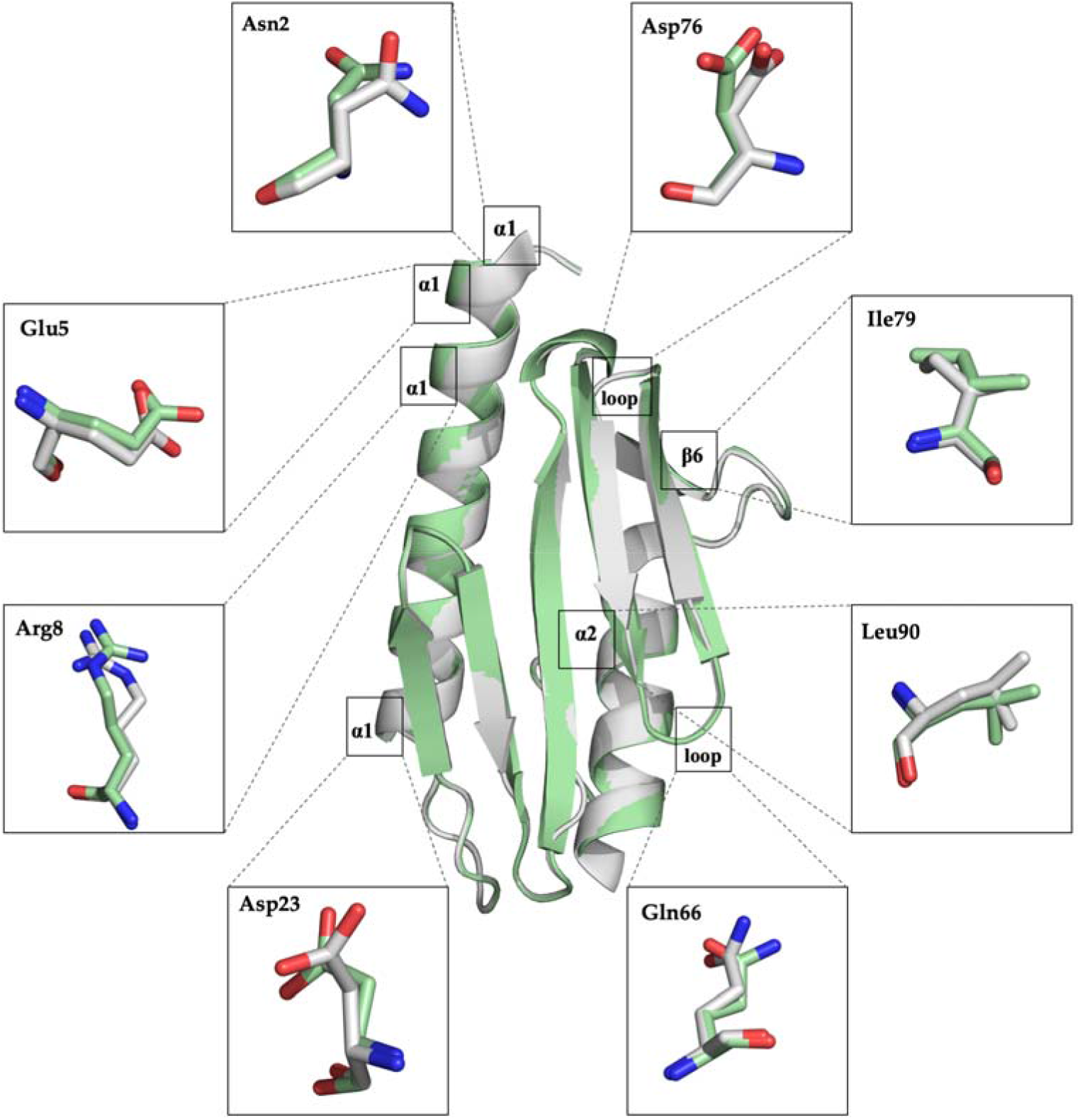
Superimposition of RT and cryogenic temperature CyaY protein. Conformational changes in side chains are depicted in boxes in stick representation..

### 3.3. Sequence Homologs and conserved residues

CyaY/Frataxin from various organisms including *Psychromonas ingrahamii37, Chaetomium thermophilum, Saccharomyces cerevisiae, Drosophila melanogaster, Homo Sapiens* (with PDB IDs: 4HS5, 6FCO, 3OFQ, 7N9I, 3S4M, respectively) were superposed with the ambient temperature CyaY structure from *E. coli* with the calculation of root mean squared deviation (RMSD) values (Figure 7). To provide a better understanding of the comparison, a multiple sequence alignment was conducted using *JALVIEW* (Figure 8). The structure-based sequence alignment analysis revealed the presence of two regions in the frataxin family proteins: a non conserved N-terminal region, which is absent in prokaryotes and imperfectly conserved in eukaryotes, and a C-terminal region, which is the functional part of the protein and exhibits sequence similarities within available homologous structures (as depicted in Figure 8-a). Highly conserved residues are notably observed in beta-sheets, particularly in β3 and β4 (Ile51, Asn52, Gln54 in β3 and Gln59, Trp61, Leu62 in β4). This finding aligns with previous reports. Other conserved residues include; Asp31 (β1), Thr40 (β2), Pro56 (3_10_), Gly68 and Asp72 (β5), Trp78 (β6), Gly84 (between β6 and *α*2), Leu91 (*α*2).

**Figure 7.**
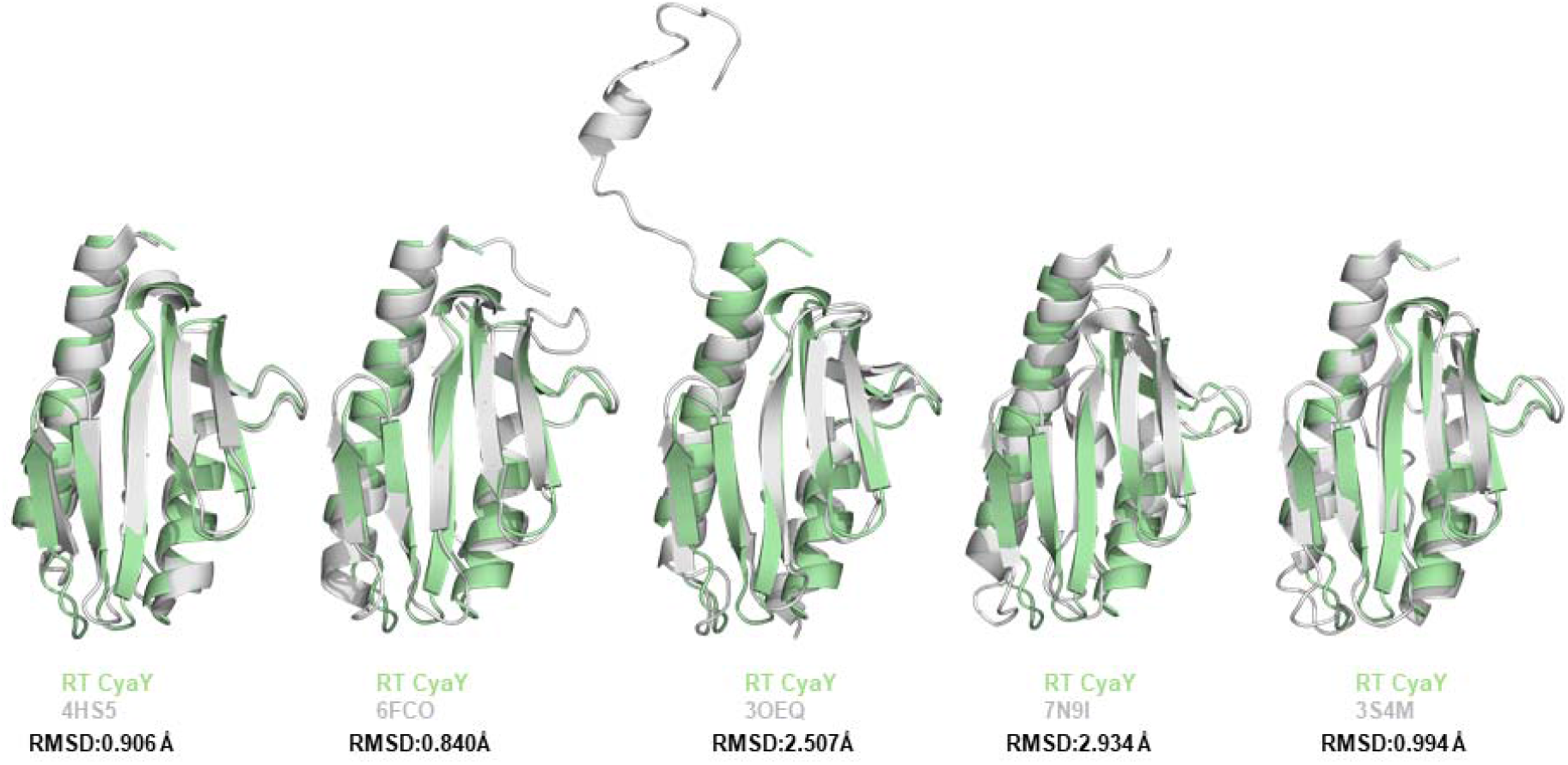
Superimposition of ambient temperature CyaY with homologous structures. Ambient temperature CyaY is colored in pale green while the superposed structure is colored in gray. Figures are generated by *PyMOL*. PDB codes of superposed structures are: 4HS5: *Psychromonas ingrahamii37* Frataxin, 6FCO: *Chaetomium thermophilum* Frataxin, 3OEQ: Yeast Frataxin, 7N9I: *Drosophila melanogaster* Frataxin, 3S4M: Human Frataxin.

**Figure 8.**
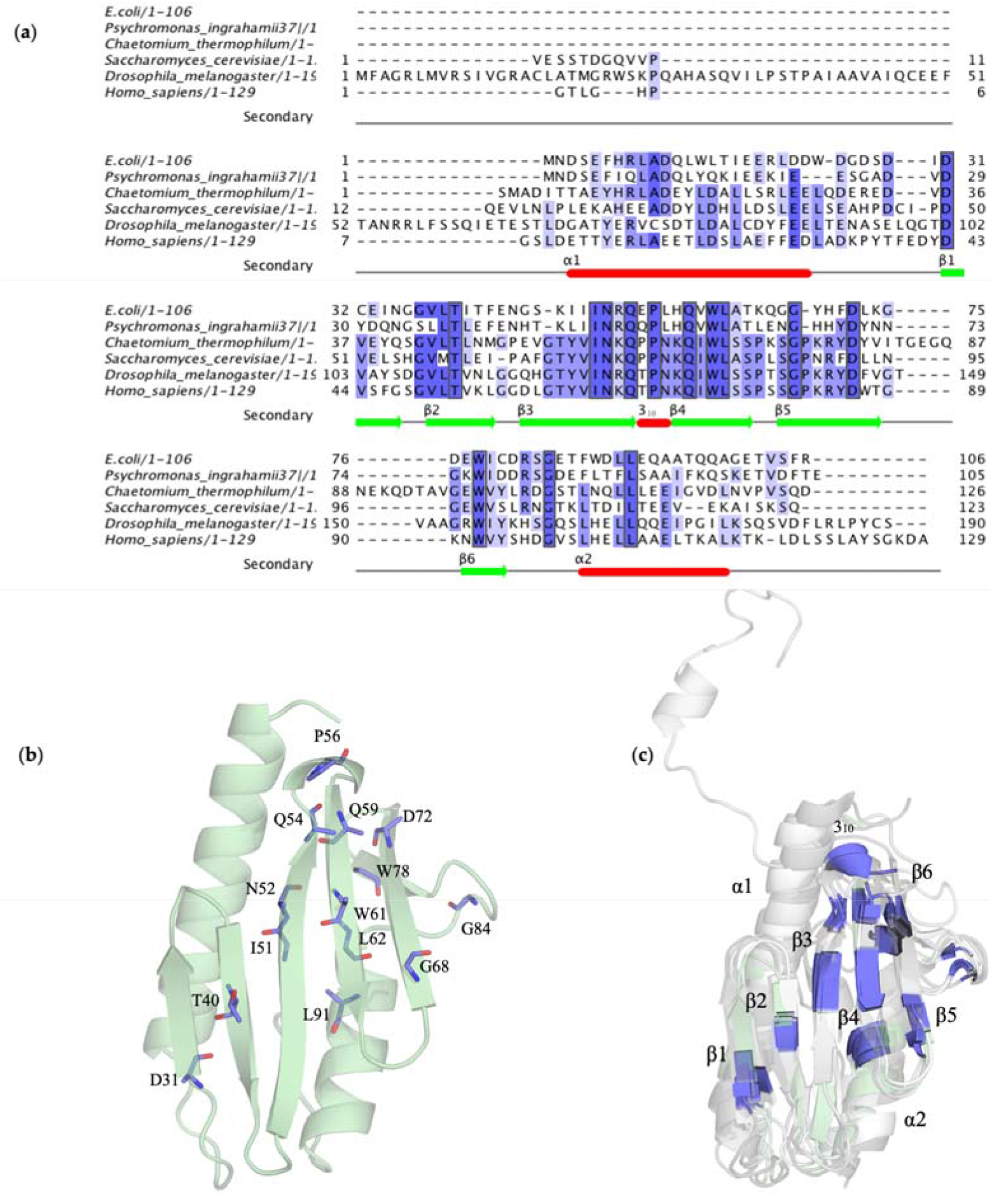
Overall representation of sequence alignment results. (a) Sequence alignment of RT CyaY from *E. coli* with 5 other members of faratxin family: PDB IDs are as following: CyaY/frataxin from *Psychromonas ingrahamii37*: 4HS5, *Chaetomium thermophilum*: 6FCO, *Saccharomyces cerevisiae*(yeasf): 3OEQ, *Drosophila melanogaster:* 7N9I, *Homo sapiens* (human): 3S4M. Black squares represent the conserved residues in the sequence alignment. (b) Conserved residues are indicated by slate color in RT CyaY structure. (c) Conserved regions are shown in all superimposed structures based on sequence alignment results in panel a.

### 3.4. Temperature factor analysis and electrostatic surface charge distribution on CyaY

In order to gain insights into the structural dynamics of the protein CyaY, we conducted a comparative analysis of its ambient and cryogenic structures using thermal ellipsoid models and electrostatic surface representations. The ellipsoid models were generated based on the temperature factors (B-factors) using PyMOL to visually highlight the differences in flexibility. Our analysis revealed that both ambient and cryogenic structures exhibited flexibility in the outer regions of the protein. However, a more pronounced intrinsic plasticity was observed in the ambient temperature structure compared to the previous cryogenic structure (Figure 9). Electrostatic forces play an established role for protein–protein interactions and protein stability. Surface charge distributions in CyaY structure in both temperatures are asymmetric, with lack of a substantial negative charge in β-sheets where there are more conserved residues (Figure 10).

**Figure 9.**
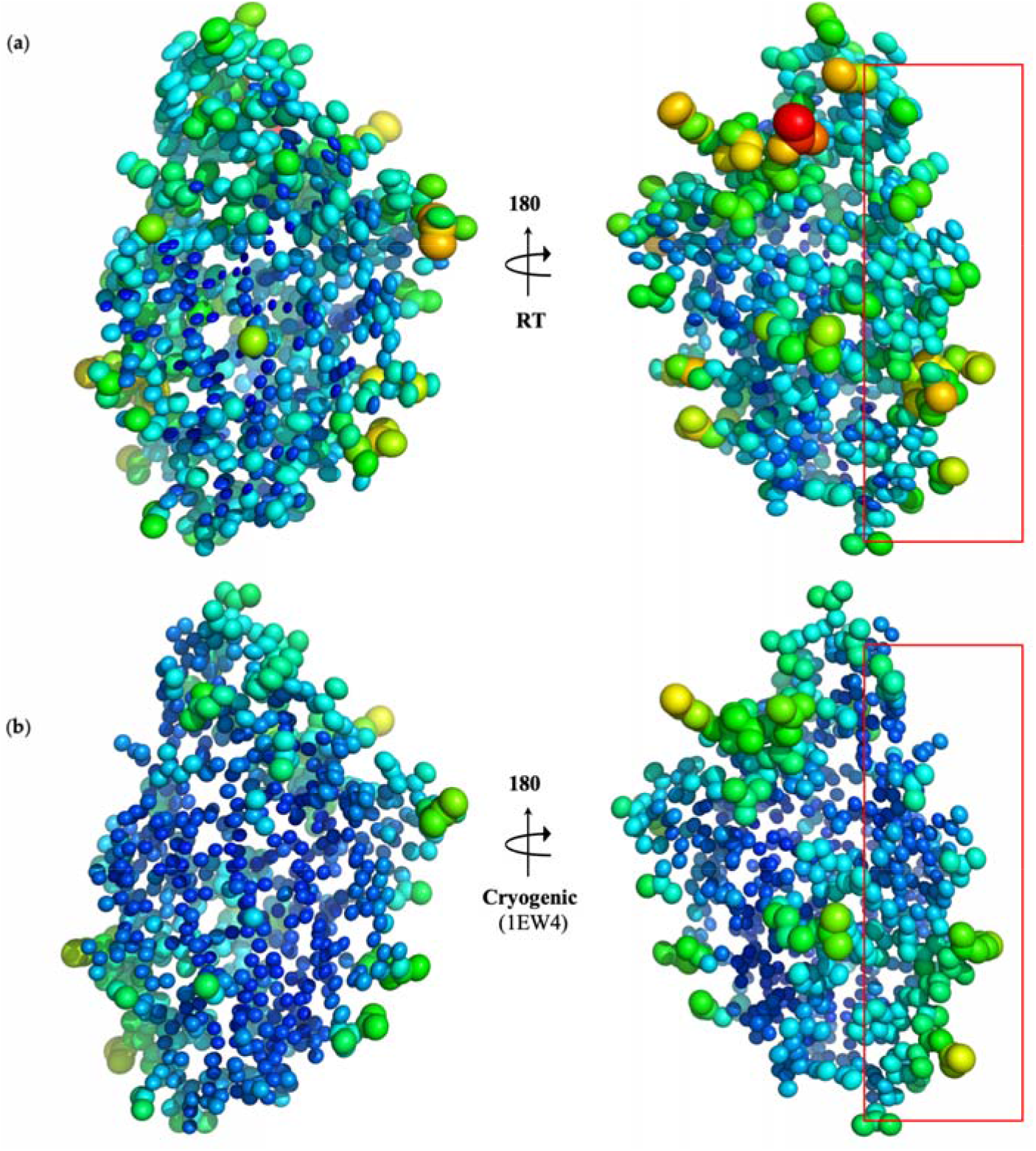
Representation of thermal ellipsoid structures of CyaY (a) ambient temperature (b) cryogenic structure. The red rectangle represents the potential binding sites on the cyaY molecule that may interact with IscS and iron. Flexible (red/orange) and stable (blue/green) regions are shown via B-factor presentation.

**Figure 10.**
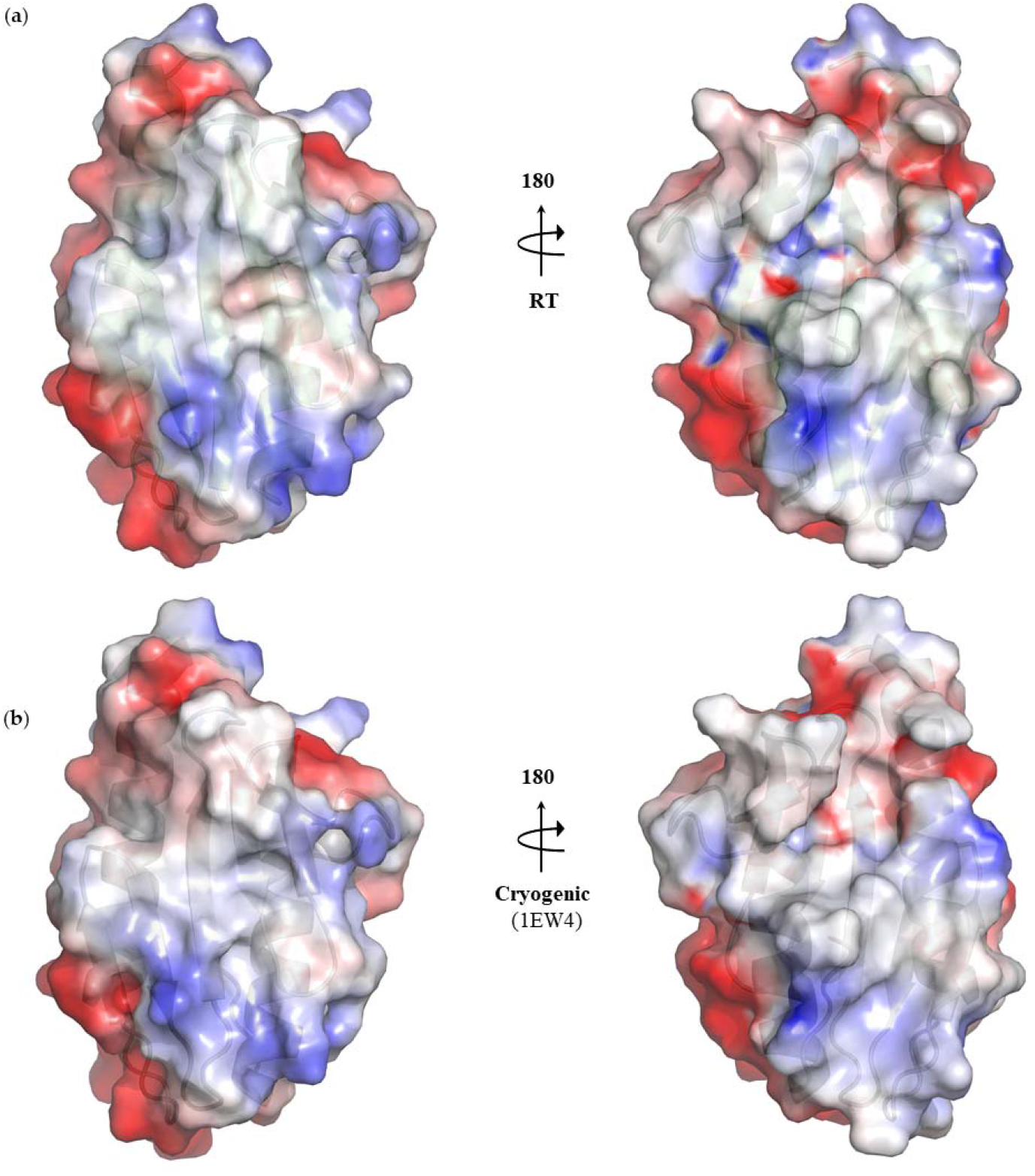
Representations of electrostatics surfaces of CyaY (a) ambient temperature (b) cryogenic structure are shown to detect the charge distribution.

## 3. Discussion

Herein, we have determined the first ambient temperature structure of CyaY from *E. coli* at 2 Å resolution. The results revealed that the ambient-CyaY structure is very similar to the previously published cryogenic structure, with a low root mean square deviation (RMSD) of 0.194 (PDB ID: 1EW4). Upon comparing the two structures, we obtained differences in their secondary structure conformations. The ambient temperature structure was found to have longer β strands compared to the cryogenic structure. This suggests that β strands may be more relaxed at ambient temperature compared to cryogenic temperatures due to the increased thermal fluctuations of side and main chains of the protein. The extended β strands may also provide protection to the hydrophobic core of the protein and enhance its stability. Our analysis also revealed variations in residues with minor conformational changes, such as Ile79 and Glu5, and significant changes in residues including Asn2, Arg8, Asp23, Asp76, Leu90, and Gln66. The observed conformational changes in various residues could impact the protein’s function, stability and interactions with its binding partners, thus, further research is required to fully understand the potential implications. It is worth noting that the ambient temperature used in our comparison was determined at a temperature which is near physiological conditions. This means that the observed differences in our structure are more close to the protein’s behavior *in vivo*.

The superposition of our ambient temperature CyaY structure with five other structures provides evidence for the presence of minor conformational differences between CyaY and its homologous structures. Despite these differences, all structures share a conserved and similar fold, indicating that the basic architecture of CyaY is well-preserved across species. Identification of conserved residues in beta-sheets and other regions of the protein, suggests that these regions are important for the structural stability and function of the protein.

We compared the flexibility of regions in both structures by visualizing ellipsoid figures generated based on the temperature factors. As expected our findings revealed the ambient temperature structure of CyaY has greater intrinsic plasticity compared to cryogenic structure. This is supported by the larger ellipsoid that indicates greater atomic motion, which corresponds to a higher B-factor. Moreover, we observed predominant blue color in the cryogenic structure, indicating higher level of rigidity. We observed a more flexible iron-binding site and IscS-binding surface for CyaY under ambient temperature conditions, these suggest the binding interactions can be via induced fit mechanism. This observation may be relevant for *in vivo* studies, as CyaY’s function in iron metabolism occurs under physiological temperatures. The absence of significant negative charge in the β-sheets, specifically in regions with highly conserved residues, shows that these residues play a crucial role in maintaining the structural integrity of the protein. Conversely, negatively charged regions of the protein tend to have interaction with other proteins and molecules.

This study sheds light on the dynamic properties of the CyaY protein under near-physiological conditions, offering a novel perspective that complements previous studies performed under cryogenic conditions. The biogenesis of iron sulfur clusters is a universally conserved complex process and not yet fully understood. There are some findings related to this topic. For example, It has been found that CyaY can bind to IscS and slow down the rates of iron sulfur cluster assembly, as a negative regulator. Additionally, it has been shown that CyaY has the potential to regulate the overall process of cluster assembly since it can transfer iron sulfur cluster from IscS to IscU. Despite these progress, still several aspects of biogenesis of iron-sulfur clusters need further investigation. For example, there are still unanswered questions regarding the precise interaction of CyaY and other key proteins, such as IscU and IscS. Our results provide a foundation for further investigation into the biogenesis of iron-sulfur clusters, including the role of key proteins such as IscS and IscU, the structural and conformational heterogeneity of complexes such as IscS-CyaY, IscS-IscU, and IscS-IscU-CyaY, and the precise transfer mechanisms of iron and sulfur in near-physiological conditions.

Understanding the mechanism of iron sulfur cluster biogenesis is crusial since any defects in this pathway may cause various human diseases, including Friedreich’s ataxia. This study paves the way for developing improved strategies for regulating iron-sulfur cluster biogenesis with potential applications in the treatment of related diseases.

## Author Contributions

Conceptualization, A.S., J.H.K.,and H.D.; methodology, A.S. and H.D.; software, A.S. and H.D.; validation, A.S., J.H.K., and H.D.; formal analysis, A.S., N.B., J.N., J.H.K., and H.D.; investigation, A.S., J.H.K., and H.D.; resources, A.S., J.H.K. and H.D.; data curation, A.S., J.H.K., and H.D.; writing—original draft preparation, A.S., J.H.K., and H.D.; writing—review and editing, A.S., N.B., J.N., J.H.K., and H.D.; visualization, A.S., J.H.K., and H.D.; supervision, H.D.; project administration, J.H.K. and H.D.; funding acquisition, J.H.K. and H.D. All authors have read and agreed to the published version of the manuscript.

## Data Availability Statement

CyaY protein structure and corresponding structure factors have been deposited into Protein Data Bank under the PDB accession code 8HZ1. Any remaining information can be obtained from the corresponding author upon request.

## Acknowledgements

Authors would like to dedicate this manuscript to the memory of Dr. Albert E. Dahlberg and Dr. Nizar Turker. The authors gratefully acknowledge the use of the services and Turkish Light Source (*Turkish DeLight*) X-ray facility at the University of Health Sciences, Experimental Medicine Application & Research Center (DETUAM), Validebag Research Park. The authors also gratefully acknowledge use of the services and facilities at Koç University Isbank Research Centre for Infectious Diseases (KUIS-CID).

